# Development of a New N-Terminomic Method to Study the Pathodegradome of the *Staphylococcus aureus* V8 Protease in Human Neutrophils

**DOI:** 10.1101/2024.11.08.622692

**Authors:** Emilee M. Mustor, Andrew M. Frey, Mary-Elizabeth Jobson, Dale Chaput, Lindsey Neil Shaw

## Abstract

*Staphylococcus aureus* is a notorious human pathogen that relies on an array of virulence factors to engender infection and evade the host-immune system. Among these are the secreted proteases, which promote pathogenesis by degrading host proteins and modulating host-defenses. Human neutrophils play a pivotal role in these defenses, acting as the first responders against invading bacteria. While many *S. aureus* effectors of virulence have been shown to target leukocytes, there is limited knowledge on how the extracellular proteases modulate neutrophil fate. Typically, protease substrates have been identified in isolated settings using one at a time approaches; with neutrophil targets few and far between. Herein, we have developed a novel N-terminomic methodology termed TAGS-CR that can facilitate global substrate characterization in streamlined manner. We thus present the application of TAGS-CR to unravelling the human neutrophil pathodegradome of the *S. aureus* V8 protease. In so doing, we captured ∼350 V8 targets, revealing critical insight into how this virulence factor can modulate neutrophil functionality on various levels relevant to *S. aureus* disease progression. We recorded cleavage of proteins necessary for neutrophil adhesion and migration, a fundamental process necessary for pathogen clearance. Furthermore, we highlight V8 cleavage of proteins involved in important neutrophil defense tactics, such as degranulation and reactive oxygen species production. This protease may also facilitate bacterial dissemination via the intentional activation of neutrophil apoptosis. Collectively, this work deepens our understanding of host-pathogen interaction and begins to unravel how *S. aureus* proteases can induce immune dysregulation through the targeting of leukocytes.

**Importance:** During infection *Staphylococcus aureus* must engage and evade the host immune system in order to successfully cause disease. As neutrophils represent the frontline of defense against invading *S. aureus* cells, it becomes increasingly important to decode how this bacterium subverts their host-defense tactics. While the contributing role to neutrophil engagement for many *S. aureus* virulence factors have been elucidated, the effects of their proteases remain largely unclear. Here, we present a novel method for global protease substrate identification, TAGS-CR, and use it to identify *S. aureus* V8 protease targets in human neutrophils. These include factors that not only govern general neutrophil function but moreover, their defense mechanisms, such as migration, degranulation, oxidative defense, phagocytosis and apoptosis.

## INTRODUCTION

*Staphylococcus aureus* is an adept human pathogen that is uniquely skilled at causing a wide array of diseases, ranging from skin and soft tissue infections, to more invasive conditions, such as osteomyelitis, endocarditis, pneumonia, and bacteremia (1). The ability to engender these various infections stems from a cadre of tightly regulated virulence factors produced by *S. aureus*, ranging from adhesins to hemolysins, toxins, lipases, proteases and other exoenzymes (2). These virulence determinants function to not only wreak havoc on host tissues but are also geared toward immune evasion and hijacking of the innate immune system (3).

In keeping with this, *S. aureus* is known to target polymorphonuclear leukocytes (PMNs or neutrophils), which serve as professional innate immune phagocytes in the host (4). PMNs are considered the primary line of defense against *S. aureus* infections in humans and are thus indispensable if pathogen clearance from the host is to be successful (5). Once at the infection site neutrophils can engulf opsonized *S. aureus* and employ mechanisms geared towards pathogen killing. Perhaps the most effective antibacterial tactic is the production of reactive oxygen species (ROS) via the NADPH oxidase complex (6,7). These molecules function to not only elicit damage directly to the bacterial membrane and intracellular components, but also enhance the immune response of PMNs (8). In addition to ROS, neutrophils also promote pathogen clearance via degranulation (9).

Consequently, neutrophil extravasation and defense present opportunities that *S. aureus* is well known to take advantage of (3). Indeed, it possesses a repertoire of virulence factors that have been shown to inhibit neutrophil activation and chemotaxis, including chemotaxis inhibitory protein (CHIPS), formyl-peptide receptor Like-1 inhibitory protein (FLIPr), and staphylococcal superantigen proteins SSL3, SSL4, SSL5 and SSL10 (10–12). In addition, it also possesses immune evasion proteins such as staphylococcal enterotoxin-like toxin X (SElX), extracellular adherence protein (Eap), and again SSL5 as well as SSL11, which are all geared towards hindering neutrophil adhesion to host epithelial surfaces (12). If neutrophils reach the site of infection, their function can be further inhibited by *S. aureus* via strategies to prevent opsonophagocytosis via complement inhibitory proteins, as well as blocking IgG antibodies bound to its surface (12). Further, if engulfed by PMNs, *S. aureus* can produce another set of immune evasion molecules to circumvent the antimicrobial activities of the phagolysosome. These include superoxide dismutases (SodM and SodA), catalase (KatA), staphylococcal peroxidase inhibitor (SPIN), Flavohemoprotein (Hmp), alkyl hydroperoxide reductase (AhpC), and staphyloxanthin, to neutralize ROS (13–18). Furthermore, staphylokinase (SAK), *O*-acetyltransferase A (OatA) and Eap will inhibit bacterial destruction by antimicrobial peptides (19–21).

Despite knowledge of how these virulence factors target immune cells, the role of the secreted proteases, and how they modulate neutrophil function, is far less well understood (22). All strains of *S. aureus* produce 4 major extracellular proteases: a metalloproteinase (Aureolysin, *aur*), a glutamyl endopeptidase (*sspA* or the V8 serine protease), and two cysteine proteases known as Staphopain A (*scpA*) and Staphopain B (*sspB*) (23). Additionally, depending on the sequence type, strains also produce up to 9 serine-protease-like enzymes known as the Spls (24). Contribution of these exoenzymes to infection is dual sided, with *S. aureus* proteases regulating the progression of infection by modulating the stability of other virulence factors, whilst at the same time having their own role in cleavage of host defense molecules (25,26). In a few instances, staphylococcal proteases have been shown to inhibit neutrophil function, for example staphopain A cleaves chemokine receptor 2 (CXCR2) resulting in diminished recruitment (27). Staphopain B also induces cleavage of CD11b and CD31, impairing neutrophil activation and survival, respectively (28,29). While V8, Staphopain B and Aureolysin were found to be produced following neutrophil phagocytosis, knowledge of their phagolysosomal targets is at best limited (30); with only the neutrophil antibacterial peptide LL-37 revealed to be a substrate (31,32).

Previously, our group deployed N-Terminomic methodologies as a global strategy to identify targets of the V8 protease in human serum (33). Approximately 90 substrates were identified in this study, with many found to be directly relevant to *S. aureus* pathogenesis, including proteins of the clotting cascade, complement system and protease inhibition networks (33). Herein, we use a newly developed N-terminomic methodology to explore the pathodegradome of the V8 protease in human neutrophils. In so doing, we were able to enhance target identification, capturing 346 V8 substrates. Our results demonstrated that the V8 protease adopts a holistic approach to neutrophil engagement and targets multiple aspects of neutrophil functionality, ranging from migration to phagocytosis, intracellular defenses and apoptosis.

## RESULTS and DISCUSSION

### Development of the TAGS-CR N-terminomic methodology

We previously published the use of TAILS N-terminomics to study the pathodegradome of the V8 protease from *Staphylococcus aureus* (33). Herein, we have developed terminal amine guanidination of substrates-charge reversal (TAGS-CR) as a novel N-terminomic methodology that uses an alternative enrichment strategy avoiding the use of single-use polymers. Our workflow is outlined in **Figure 1** and uses a combination of previously defined chemistries for the identification of protease substrates (34,35). Guanidination (Gd) of lysine ε-amines and N-terminal α-amines was our chosen labelling strategy as this method not only negates digestion inefficiency seen with more traditional approaches, such as dimethylation or acetylation modifications as a result of blocked lysine residues, but also increases ionization efficiency of peptides (34). Gd-labeled samples undergo reduction, alkylation, and tryptic digest prior to downstream enrichment via charge reversal (CR) of internal peptides and strong cation exchange (SCX) (35,36). Post digest, peptides undergo a series of incubation steps whereby internal, unlabeled tryptic peptides gain a negative charge via sulfation. Strong cation exchange then facilitates enrichment of Gd-labelled peptides, which had amines already converted to guanidinyl groups prior to tryptic digest. Thus, the native- or neo- (i.e. caused by proteolysis prior to trypsinization) N-termini are retained during SCX, but negatively charged tryptic peptides can no longer bind to the SCX matrix, thus facilitating their removal.

**Figure 1.**
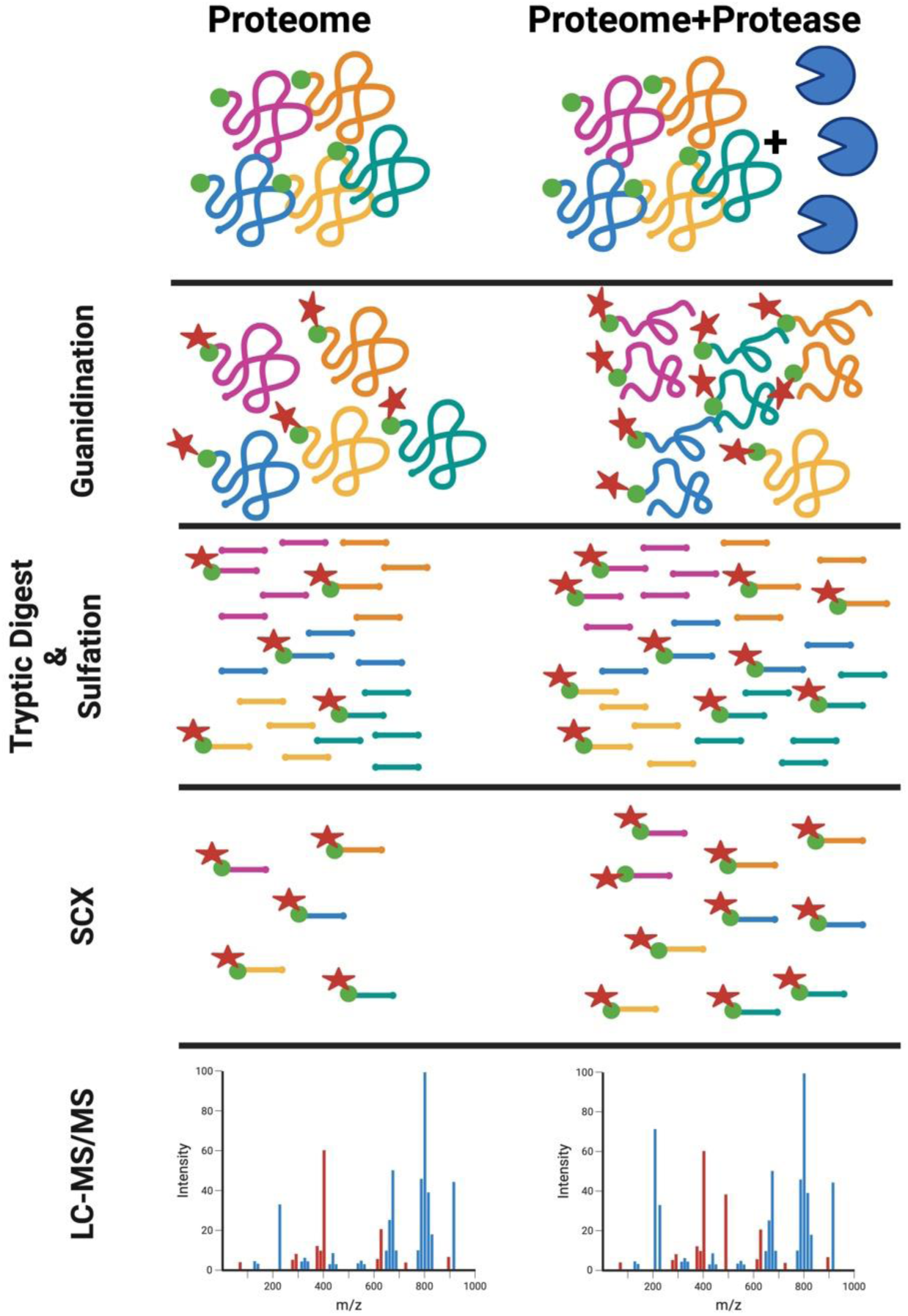
Guanidination and Enrichment of N-terminal peptides through TAGS-CR. Proteins are reduced and cysteine residues are blocked via alkylation prior to guanidination of N-terminal α-amines and lysine ε-amines (not shown) using 1*H*-Pyrazole-1-carboxamidine (HPCA). A manufacturer provided S-trap protocol is used and N-terminally labelled proteins are tryptically digested. Following elution, peptides undergo (di)sulfonation of free amines that have been liberated post tryptic digest. Strong cation exchange (SCX) is then performed to enrich for both native and neo-N-termini, the latter of which indicates a cleavage event and thus only pertains to the protease treated condition. LC-MS/MS analysis is then performed on the TAGS-CR enriched samples to obtain the N-terminome.

### Uncovering V8 substrates within neutrophil proteomes

Utilizing TAGS-CR, we sought to expand our studies with the V8 protease of *S. aureus*, deploying our new method to explore the pathodegradome of human neutrophils. Accordingly, the proteomes of differentiated HL-60 cells were treated with V8 protease and, employing a binary comparison of N-termini between treated and untreated samples, TAGS-CR identified 346 V8 protein substrates (**Table S1**). Of these, we captured 454 cleavage events unique to the V8 treated condition (**Table S2**). It is well established that the V8 protease cleaves primarily at the carboxyl side of glutamic acid (E) and to a lesser extent aspartic acid (D) residues (37). As such, we sought to initially validate our V8 targets via analysis of the amino acids surrounding the identified cleavage sites (**Figure 2A and 2B**). Of the 454 neo-N-termini, glutamic acid (E) accounted for ∼90% of the residues upstream of the scissile bond at position P1 (**Figure 2A**). Sequence analysis with ICELOGO (38) was also used to assess sequence conservation surrounding the cleavage position (P5-P5’) (**Figure 2B**). In keeping with the specificity of the V8 protease, we found that only glutamic acid was significantly increased at P1 (**Figure 2B**). We also observed a significant increase in alanine and glycine residues at P1’, leucine residues at position P2 and valine residues at P4’ (**Figure 2B**). Our previous study of V8 targets in human serum revealed a similar trend in amino acids at these positions, indicating a likely consensus sequence for V8 protease substrate preference (33). Additionally, in our past (33) and present study we noted an apparent inhibitory effect of proline on the activity of the V8 enzyme due to decreased frequency at positions P3 and P1’. The inhibitory effect of proline on not only V8 activity but protease activity in general has previously been recorded in the MEROPS database, and in a number of *in vitro* experiments (39–41). The similarity in cleavage consensus sequence between our study and those previous, indicates that changing methods from TAILS to TAGS-CR did not bias identification of cleavage sites, offering additional validation of TAGS-CR, while providing a new and diverse set of host substrates for V8.

**Figure 2.**
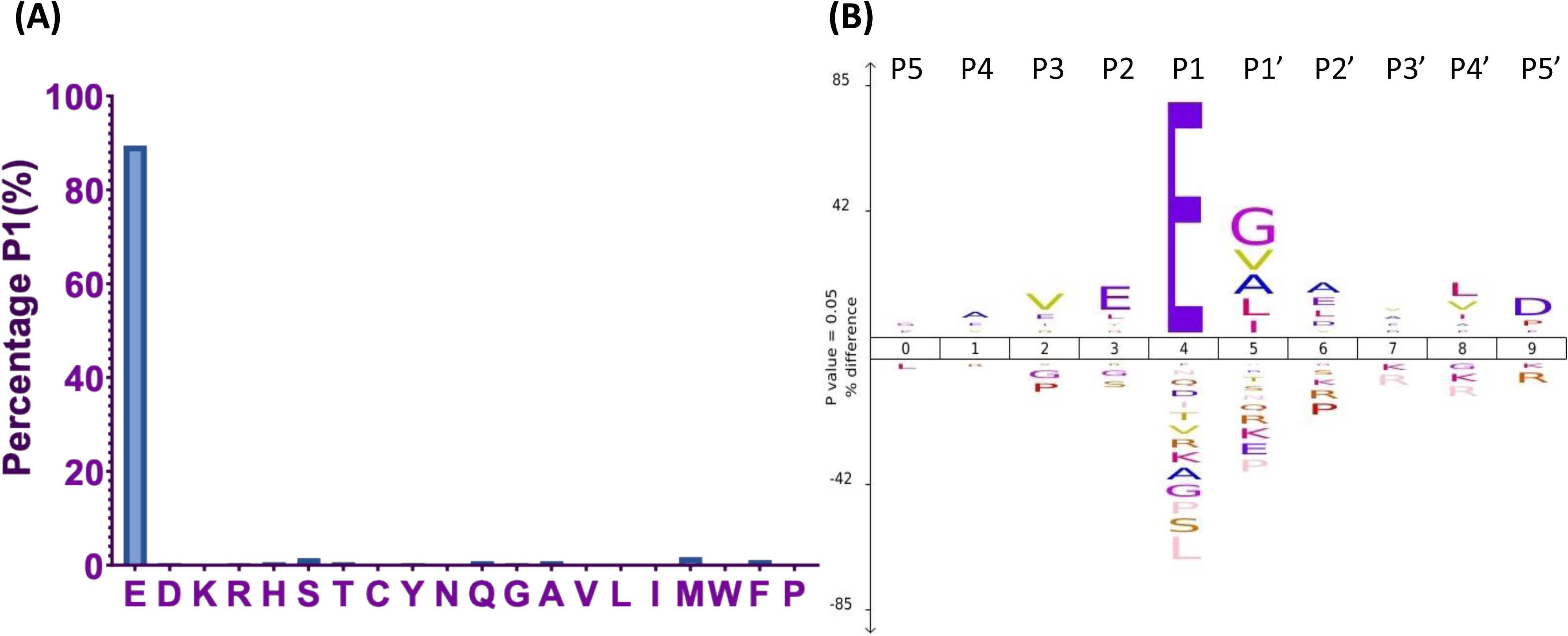
TAGS-CR validates cleavage specificity of the V8 protease. Analysis of the amino acid sequence surrounding the site of proteolysis provided immediate validation of V8 induced cleavage of our 346 neutrophil targets by showcasing the known precision of this enzyme to cleave primarily at the carboxyl side of glutamic acid (E) residues. Demonstrating this, ∼90% of residues upstream of the cleavage site accounted for glutamic acid (E) residues (**A**). We performed an additional dimension of analysis using ICELOGO whereby we looked at the percentage difference of significantly increased and decreased residues at the five positions (P5-P5’) surrounding the sessile bond. Here we found that only glutamic acid was significantly increased at P1 or upstream of the cleavage site (**B**).

### Ingenuity pathway analysis highlights host canonical signaling pathways targeted by V8

Using the Ingenuity Pathway Analysis (IPA) software, we were able to highlight canonical host signaling pathways to which V8 leukocyte targets belonged. Here, we identified 920 signaling cascades potentially being modulated by the *S. aureus* V8 protease (**Table S3**). Among these included, leukocyte extravasation signaling, integrin cell surface interactions, neutrophil degranulation, Rac2 signaling and other oxygen-dependent ROS production pathways, phagocytosis and phagosomal maturation, and finally, the intrinsic pathway for apoptosis (**Table S3 and Figure 3**). This analysis suggests that the V8 protease can hinder neutrophil functionality on many different levels relevant to *S. aureus* pathogenesis.

**Figure 3.**
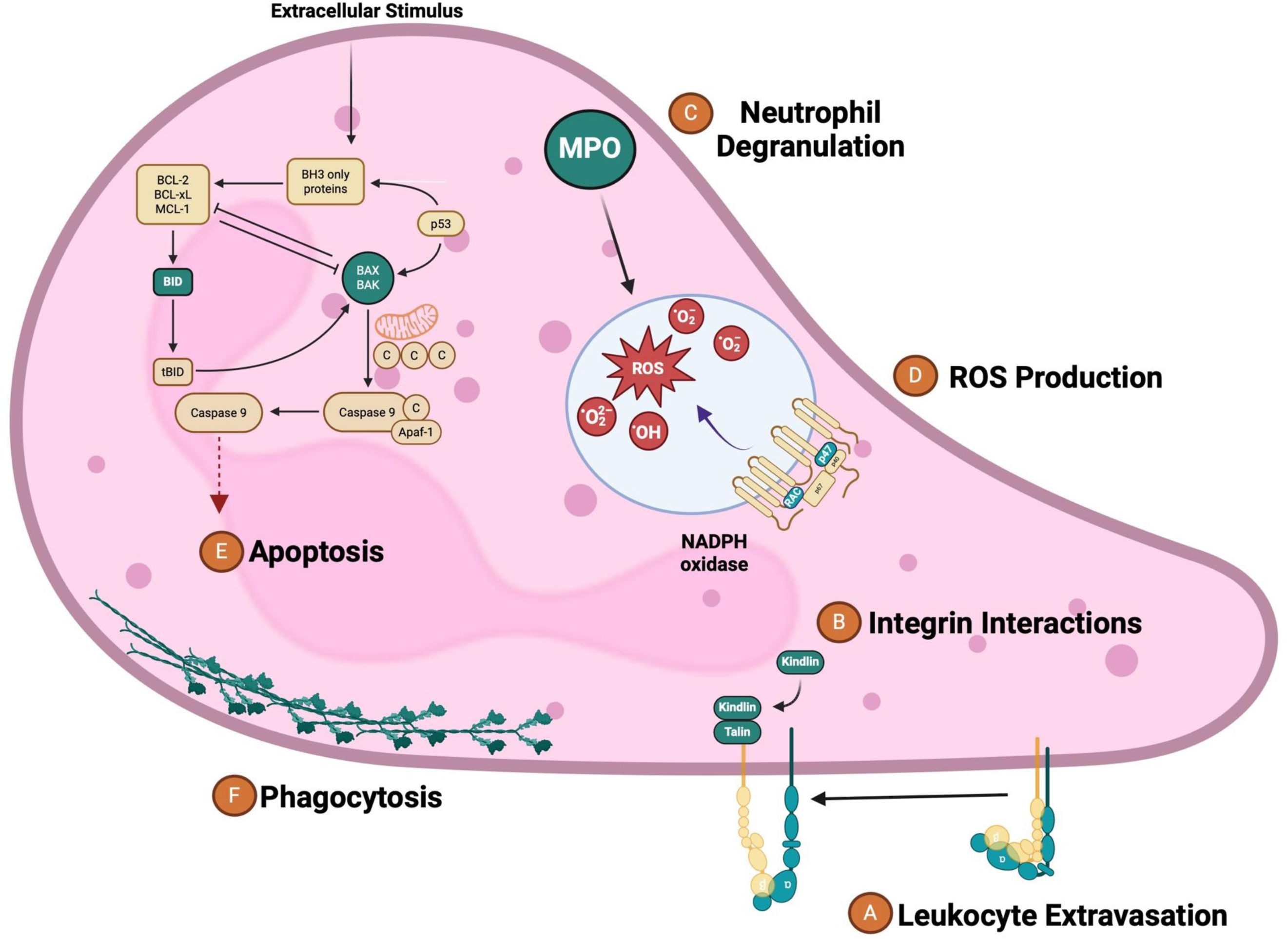
V8 targets multiple dimensions of neutrophil functionality and defense relevant to *S. aureus* pathogenesis. Green molecules are highlighted as key neutrophil proteins cleaved by V8. Displayed adjacent to these are the corresponding neutrophil processes they play a role in. These include: (**A**) Leukocyte Extravasation, (**B**) Integrin Interactions, (**C**) Neutrophil Degranulation, (**D**) ROS Production, (**E**) Apoptosis, and (**F**)Phagocytosis.

### TAGS-CR highlights V8 targets essential for leukocyte extravasation

Neutrophil recruitment is an integral process typically involving various receptors on the surface of cells (42). Once neutrophils become activated, they adhere and migrate through host tissues to different sites of infection via transcellular or paracellular routes, in a process termed neutrophil extravasation (42). TAGS-CR captured V8 induced cleavage of various receptor proteins belonging to this signaling pathway, such as intracellular adhesion molecule 3 (ICAM3), guanine nucleotide-binding protein G(i) subunit alpha-2 (GNAI2), transforming protein RhoA (RHOA), Ras-related C3 botulinum toxin substrate 2 (RAC2), moesin (MSN), and integrin alpha-L (ITGAL). Upon closer assessment, the primary influence of V8 on this process is seemingly geared towards modulating the adhesion dynamics of neutrophils. For example, moesin belongs to a group of adaptor proteins, collectively termed ERM (ezrin/radixin/moesin), that influence leukocyte response (43). Previous studies have demonstrated moesin and moesin/ezrin knockout neutrophils present with abrogated adhesive abilities (44). V8 could potentially mimic this effect as cleavage, occurring at position 396 in the alpha helix domain could diminish functional activity of moesin. In its active and subsequently unfolded state, this protein acts as a bridge between actin filaments and the plasma membrane via their interaction with opposing ends of this molecule (43). While this protein’s FERM domain mediates attachment to the plasma membrane, cleavage at any position would destroy the linkage effect of moesin, resulting in dissociation of actin and plasma membrane structures with the overall effect of potentially hindering neutrophil adhesion. Similarly, we also captured Rac2 and RhoA as V8 substrates. These proteins belong to the Rho family of small GTPases, and, while they are multifunctional, regulating numerous neutrophil activities, importantly they also control neutrophil adhesion (45,46). Moreover, this process of neutrophil arrest could further be modulated by V8 cleavage of ICAM3 and ITGAL, both of which facilitate neutrophil adherence to epithelial surfaces via binding to their respective ligands (47).

Integrins, like ITGAL, have been found to play pivotal roles in promoting not only neutrophil adhesion and movement but also effector functions during inflammatory conditions such as *S. aureus* infection (48). V8 cleaves ITGAL (CD11a) at positions 477 and 510, located in the FG-GAP-5 and FG-GAP-6 repeats, respectively. The N-terminal region of the integrin alpha subunit contains seven FG-GAP repeats, of about 60aa each, that fold to form a beta propellor domain important for ligand binding (49). While both magnesium and calcium ions bind this beta propellor domain and activate the integrin alpha subunit, V8 activity would disrupt calcium binding as cleavage of this molecule occurs in the middle of a calcium binding motif present in the FG-GAP-6 repeat (Protein Data Bank in Europe Knowledge Base, PDBe-KB) (49). While TAGS-CR only captured cleavage of the integrin alpha subunit, CD11a, these molecules are typically heterodimeric in form (50). As such, the ITGAL chain (CD11a) will combine with the integrin beta 2 chain (ITGB2 or CD18) to form the leukocyte function-associated antigen (LFA)-1, also known as a β2 integrin, a complex integral to neutrophil migration upon ligand binding (50). Highly expressed on the surface of neutrophils once activated by cytokines, LFA-1 will bind to ICAM-1 found on the surface of endothelial cells, facilitating adhesion prior to migration (50). V8 mediated cleavage of the ITGAL chain could work to dismantle this complex, perpetuating *S. aureus* survival by hindering bacterial clearance due to an insufficient number of neutrophils at the site of infection. The (LFA)-1 complex has long been a molecule of interest as defects in its expression cause leukocyte adhesion deficiency (LAD)-1 syndrome in the host (50). In patients suffering with LAD-1, impairment of the adhesive ability of leukocytes has been linked to mutations in the CD18 subunit of the complex resulting in nonfunctional LFA-1 (50,51). Importantly, patients suffering with LAD-1 syndrome experience recurrent bacterial infections, along with detrimental health effects such as painful lesions that fail to heal (50,51). Recurrent bacterial infections are a hallmark of *S. aureus* disease and are often seen in patients with leukocyte defects (52). *S. aureus* proteases may therefore contribute to the occurrence of these continuing infections via cleavage of the ITGAL chain by V8. This may compromise the activity of the LFA-1 complex, leading to a dampened neutrophil response during infection.

An important consideration with proteomic studies such as ours is that rigorous validation of targets must be undertaken. As such, we confirmed V8 cleavage of ITGAL via immunoblot (**Figure 4A**). In untreated neutrophil proteomes, we observed a ∼129kDa band corresponding to the predicted molecular weight of this protein, as well as an additional band accounting for a glycosylated form. Proteolytic cleavage of ITGAL by V8 at residue 510 would give rise to a C-terminal fragment ∼72kDa in size – which was observed in the immunoblot. (**Figure 4A**).

**Figure 4.**
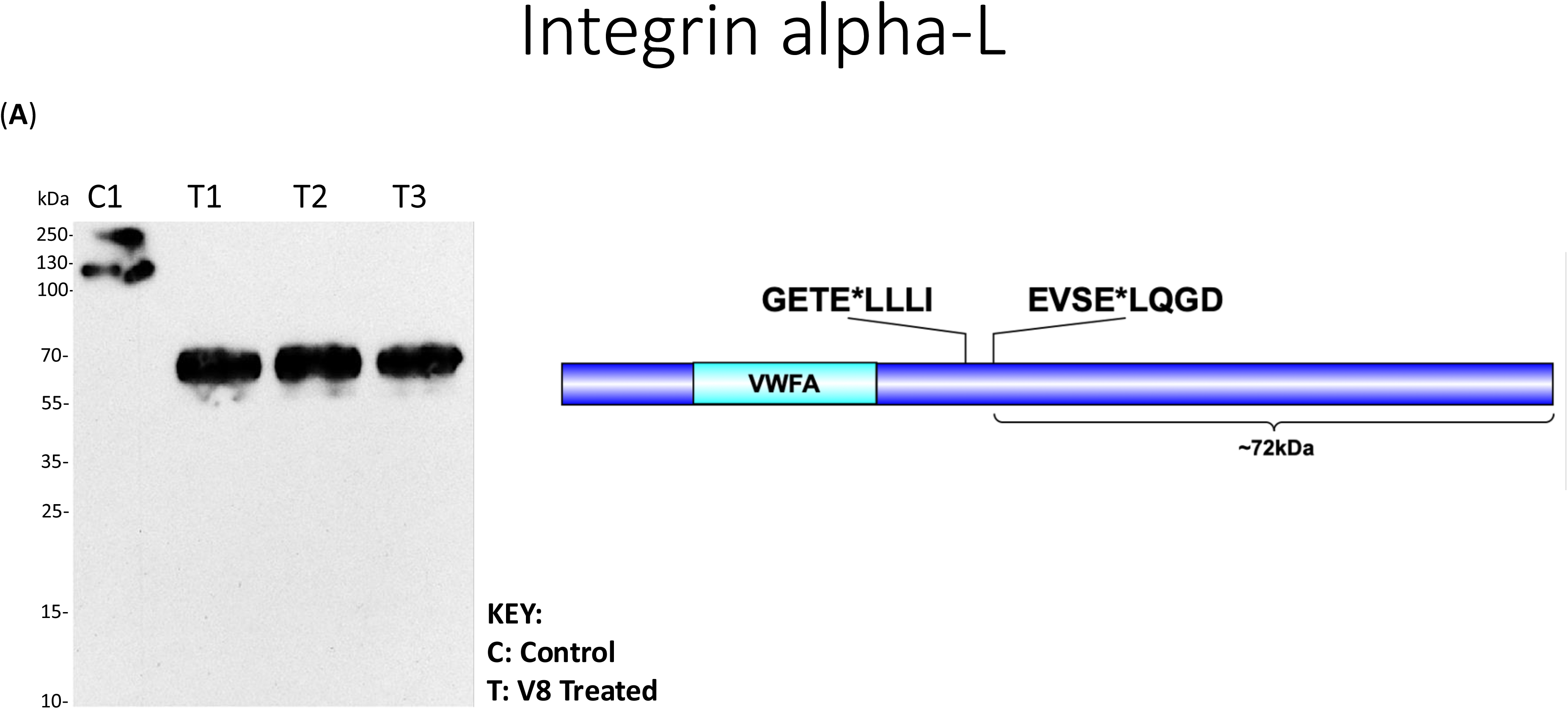
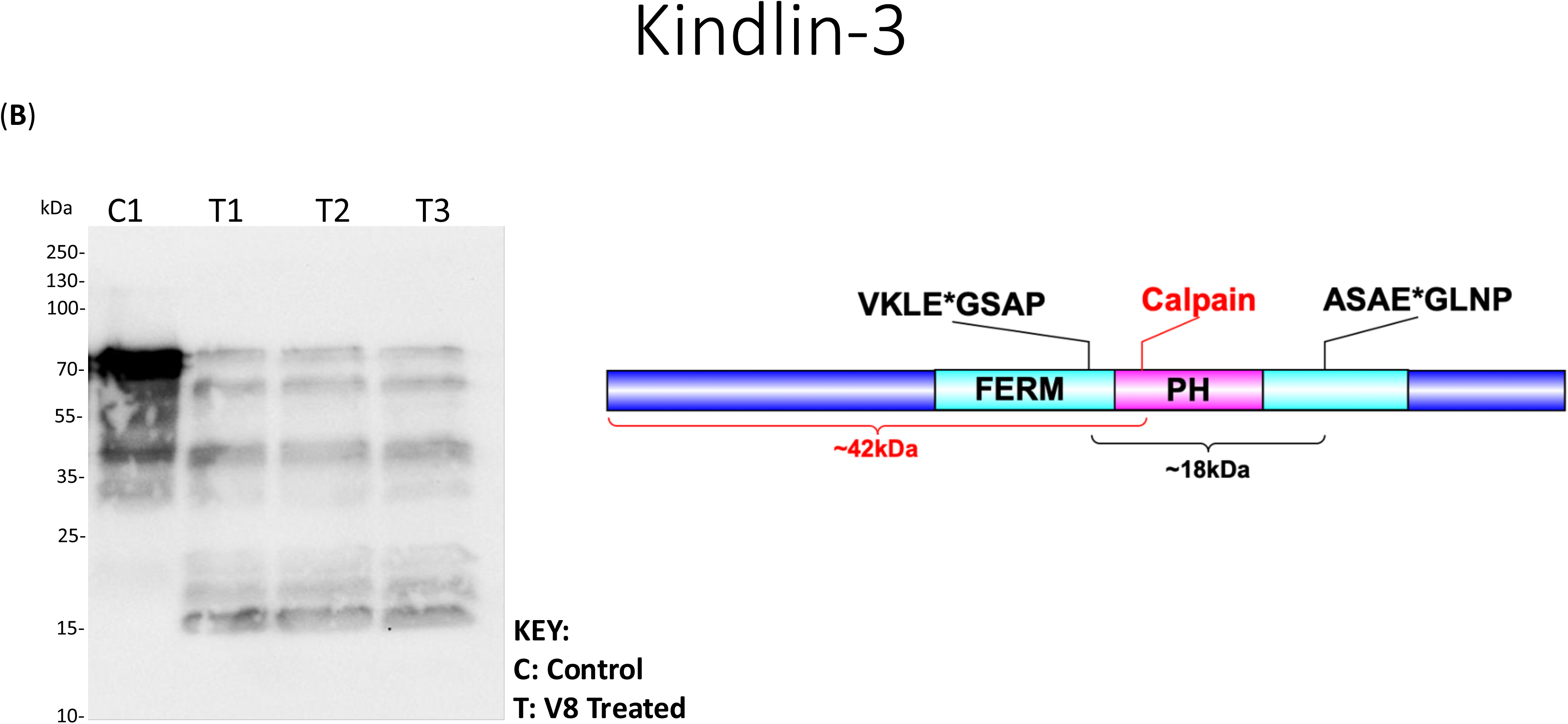
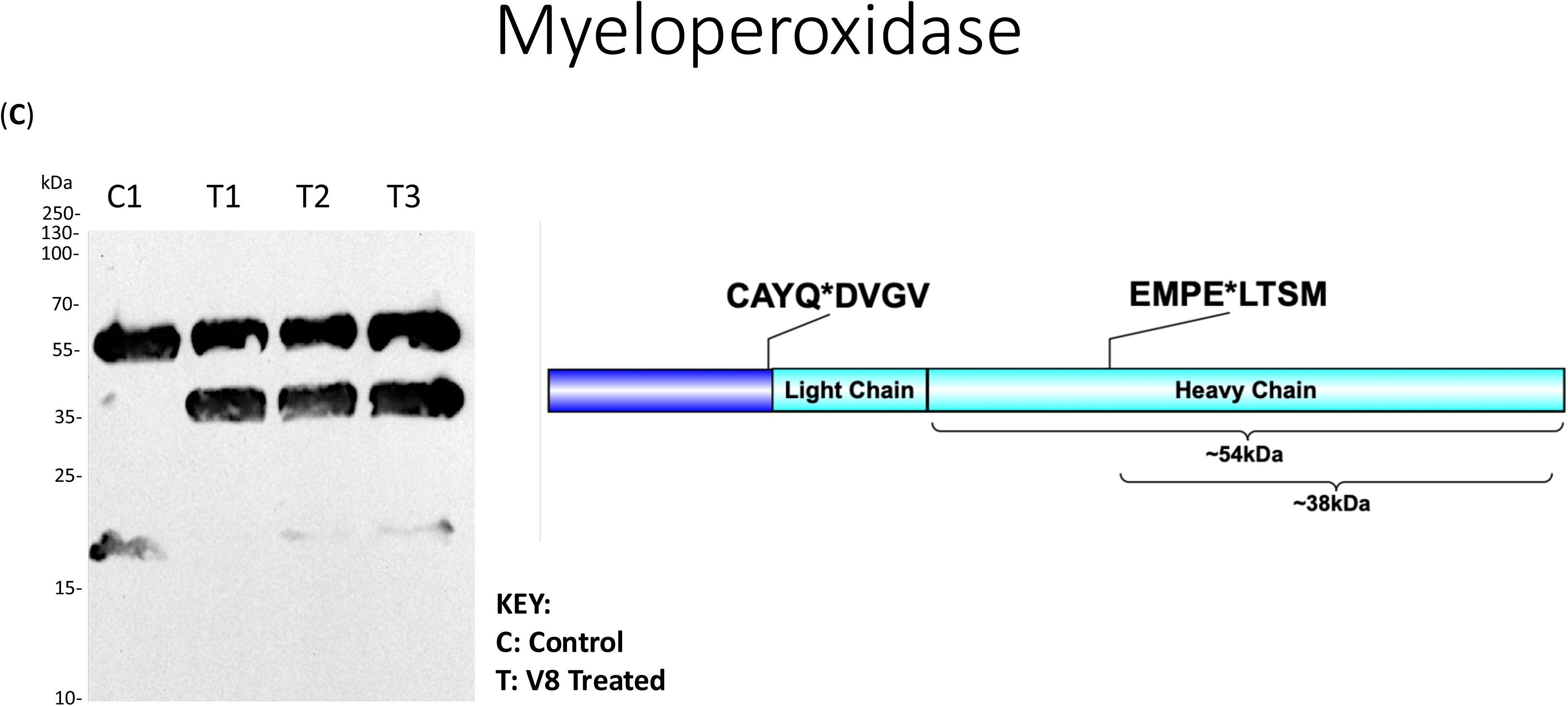
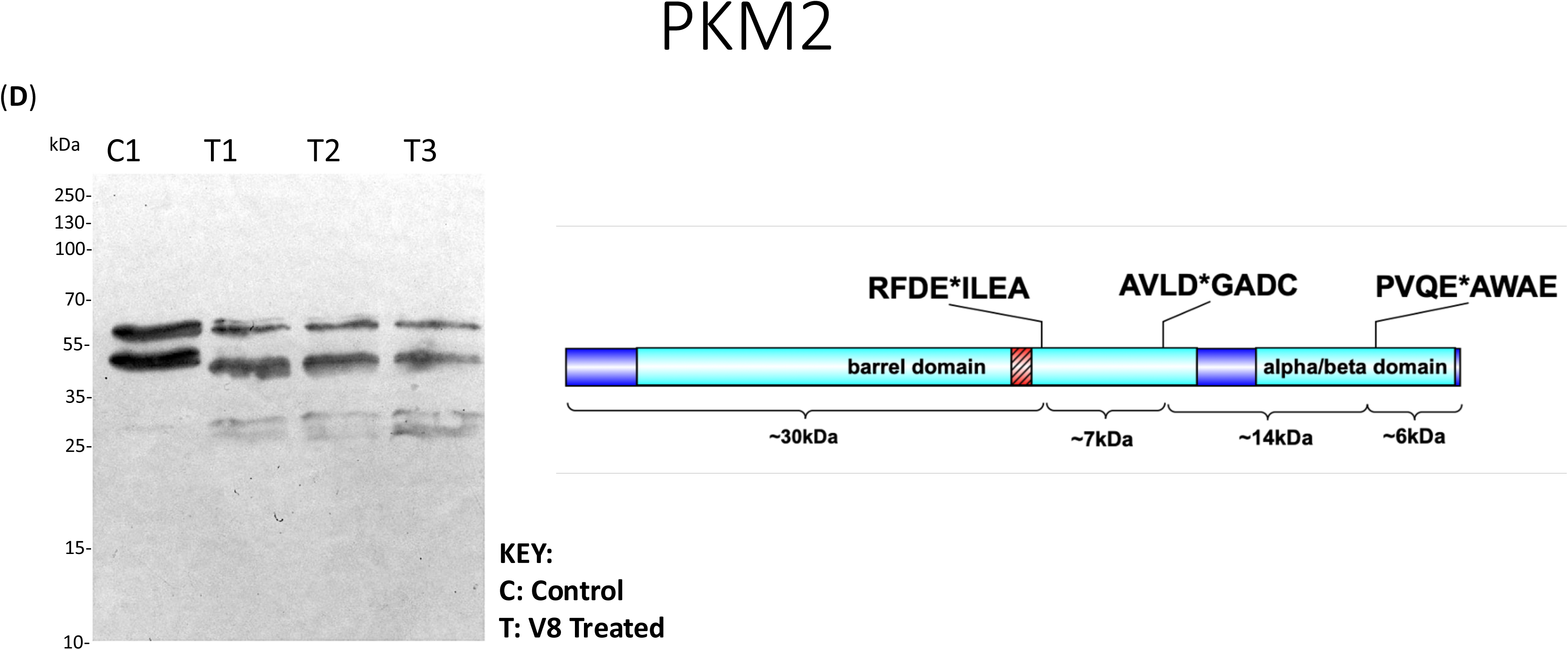
Immunoblot validation of V8 neutrophil targets. Neutrophil proteomes were exposed to 200ng of the V8 protease for 16h at 37°C whilst control conditions remained untreated. Degradation patterns and their corresponding approximate molecular weights (calculated using Bioinformatics.org) according to the cleavage events captured by our N-terminomic data are outlined in the relevant protein schematic. Also highlighted in the schematics are the identified sites of V8 cleavage. Western blot analysis of neutrophil V8 targets include: (**A**) Integrin alpha-L, (**B**) Kindlin, (**C**) Myeloperoxidase, and (**D**) Pyruvate kinase PKM2. VWFA = von Willebrand A. FERM = Ferm domain. PH = PH domain.

### V8 further hampers neutrophil migration by targeting integrin activators

Leukocyte defects come in many forms and range from type I, described above, to type III (53). While mutations in the *ITGB2* gene cause LAD type I, type III is caused by mutations in the *FERMT3* gene (54,55). Patients suffering with LAD III similarly experience recurrent bacterial infections along with other symptoms including skin and soft tissue lesions and uncontrollable bleeding (54,55). *FERMT3* encodes the fermitin family homolog 3 (Kindlin-3) protein product, which was found to be cleaved by the V8 protease (**Table S1 and S2**). In tandem with talin-1, notably also a V8 substrate, kindlin-3 is responsible for ‘inside out’ signaling to activate the LFA-1 integrin complex (56). This results in a conformational change, ultimately allowing this molecule to interact with its binding partners and arrest to the vascular endothelium prior to slow rolling along the surface and subsequent transmigration (53,56). Previous studies have described *in vivo* data where neutrophil deficiency in kindlin-3 resulted in attenuation of neutrophil arrest (56). In our study, we hypothesize that V8 cleavage of this molecule could have similar effects. Two V8 cleavage events were detected in Kindlin-3, with start positions 336 and 500, both corresponding to the FERM domain. Similar to ITGAL, kindlin-3 combines with a beta integrin subunit, in this case ITGB1, ITGB2 or ITGB3 (54). As such, the FERM domain of kindlin-3 induces activation of the beta subunit and complex formation (54); thus, V8 cleavage would disrupt this interaction. Interestingly, in patients with the LAD-III deficiency, a *FERMT3* gene mutation was seen corresponding to position 552 in the protein product, also found in the FERM domain (54). It was hypothesized that this mutation would hinder the binding capabilities of the FERM domain to its beta integrin interaction partner (54). While mutations are not akin to cleavage, our data highlights multiple avenues by which neutrophil arrest and subsequent migration may be targeted by the *S. aureus* V8 protease.

During the validation of cleavage for this pathway, immunoblotting with antibodies against kindlin-3 revealed a band corresponding to the full-length protein at ∼75kDa in both V8 treated and untreated proteomes, however with a higher intensity in the latter, as expected since no enzyme was present (**Figure 4B**). Studies have shown this protein to be heavily phosphorylated at approximately 30 sites (57), thus possibly accounting for the additional band directly below the full-length protein (**Figure 4B**). Our immunoblot also presented an unexpected banding pattern between ∼30-50kDa in both V8 treated and untreated conditions, possibly indicating that native proteolysis of kindlin-3 is occurring in the neutrophil proteome, independent of V8 treatment (**Figure 4B**). In line with this, kindlin-3 has been shown to be cleaved by calpain, an intracellular cysteine protease found abundantly in the HL60 promyelocytic leukemia cell line from which our neutrophil cells were derived (57–60). Calpain cleavage of kindlin-3, found at position 373, would yield two fragments, one of which would be ∼42kDa - directly corresponding to the fragment observed (**Figure 4B**). However, independent to calpain activity, we were still able to validate V8 cleavage of kindlin-3 at positions 336 and 500. In treated samples, we observed a cluster of 3 unique bands, one of which directly aligns with the V8 liberated N-terminal fragment at ∼18kDa outlined in the protein schematic (**Figure 4B**). Likely, the surrounding bands at ∼15kDa and ∼23kDa may correspond to modified versions of the specific V8 liberated fragment.

### Dysregulation of neutrophil degranulation by V8

Despite the many efforts of pathogens like *S. aureus* to prevent neutrophil migration, these workhorses of immunity can still reach the site of infection to engulf and clear invading bacteria from the host (61). To facilitate intracellular killing, neutrophils undergo degranulation and will generate and release potent microbicidal molecules including superoxides, other reactive oxygen species (ROS), as well as degradative enzymes and peptides (22,52,67). IPA highlighted 41 proteins involved in the process of neutrophil degranulation, as targets of V8 (**Table S3**). Upon further analysis we found several of these proteins to be directly implicated in pathogen mediated killing, either serving as an effector molecule or their precursor. Among them were cathepsin C, a degradative lysosomal cysteine peptidase known to promote pathogen clearance whilst also serving as an activator of other important neutrophil serine proteases, including elastase, proteinase 3 and cathepsin G, all constituting as azurophilic granules that exhibit a similar effect (62). Thus, our data suggests that cleavage of this molecule by V8, and therefore a potential dampening of its activity, may compromise the host’s ability to defend against *S. aureus*. This notion is further supported by previous research revealing that neutrophils lacking olfactomedin 4, a negative regulator of cathepsin C activity, demonstrated increased intracellular killing of *S. aureus* (62). Moreover, *in vivo*, mice carrying these mutated immune cells experienced enhanced clearance of *S. aureus*, largely owing to an increase in activity of this lysosomal exopeptidase, highlighting its essential role in innate immune defense (62).

Beyond cathepsin C, TAGS-CR also captured V8 cleavage of the endosomal adaptor protein p14 (LAMTOR2). Immunodeficiency in this protein has been shown to culminate in congenital neutropenia, altered microarchitecture of azurophilic granules, and importantly decreased ability for pathogen clearance within the phagosome (63). Whilst V8’s negative impact on the azurophilic granules of human leukocytes may be indirect through the targeting of cathepsin C and the endosomal adaptor protein p14, we also observed direct degradation of these key molecules via cleavage of myeloperoxidase (MPO). Myeloperoxidase is a heme driven antimicrobial enzyme that facilitates catalysis of hydrogen peroxide to hypochlorous acid, a reactive oxygen species (ROS) (64,65). Participating in oxidative defense, this potent ROS causes irreparable damage to invading *S. aureus* (66). We observe V8 cleaving MPO at position 412, downstream of its heme binding cluster at residues 495, 498-502 and 505 (PDBe-KB). Given the ferric form of MPO is required to drive the catalytic cycle of this enzyme (65), V8 cleavage near these residues would act to destabilize this molecule and decrease HOCl production. To emphasize how important MPO inhibition is for *S. aureus* pathogenesis, this bacterium encodes an additional secreted virulence factor, staphylococcal peroxidase inhibitor or SPIN, that is a specific inhibitor of MPO activity (66). As such, in addition to SPIN, our findings suggest an alternative avenue of protection for *S. aureus* against MPO-dependent killing elicited by V8. Collectively, our data suggests that V8 may assist in mitigating bactericidal activities of the neutrophil phagosome, potentially allowing *S. aureus* to survive intracellularly and further disseminate to different niches within the host.

In support of this, we present immunoblot validation of myeloperoxidase cleavage (**Figure 4**). MPO is a homodimer, with each identical monomer having a heavy chain (60kDa) and a light chain (15kDa) (68,69). In the untreated control, immunoblotting revealed the heavy chain of myeloperoxidase at ∼60kDa (**Figure 4C**). In V8 treated proteomes, we still observed the presence of the heavy chain, however, an additional, major cleavage product appeared at ∼38kDa (**Figure 4C**). This corresponds to the C-terminal fragment of myeloperoxidase produced following V8 cleavage of the heavy chain at position 412 (**Figure 4C**).

### Oxygen-dependent host-defense mechanisms are vulnerable to V8 activity

Continued targeting of ROS production by V8 was observed via cleavage of multiple proteins involved in activation of the NADPH or phagocyte oxidase complex that governs respiratory burst and superoxide production within neutrophil phagolysosomes (6). Neo-N-termini arising from V8 treatment were found belonging to not only downstream enzymes modulating the activity of the complex, such as pyruvate kinase M2 (PKM2), protein kinase C alpha type (PKC-alpha), and Ras-related C3 botulinum toxin substrate 2 (Rac2), but also a physical subunit of the NADPH oxidase complex, putative neutrophil cytosol factor 1C (p47^phox^ or NCF-1C) (70–73). NADPH oxidase activity is ultimately dependent on glycolysis and glycolytic intermediates to trigger the assembly of its subunits and subsequent ROS production (70). Herein, we see a potential inhibitory effect of the V8 protease on this process as PKM2 is the rate limiting enzyme, and once allosterically activated, controls ROS production via mediation of glycolytic intermediates (70). Emphasizing the importance of this molecule to *S. aureus* pathogenesis, deletion of PKM2, or its pharmacological inhibition, results in decreased killing of *S. aureus in vitro* as a consequence of ablated ROS production (70). Furthermore, *in vivo, Pkm2* deficient mice infected with *S. aureus* show increased bacterial loads and delayed wound healing at the site of infection (70). Products of glycolysis regulated by PKM2, namely diacylglycerol or DAG, will go on to activate PKC-alpha in a calcium dependent fashion (70). This isozyme then phosphorylates the p47^phox^ subunit of the NADPH complex, in turn leading to its activation and ROS production (70–71,73). V8 was found to cleave both PKC-alpha and p47^phox^ at position 283 and position 33, respectively. While PKC-alpha is cleaved by V8 directly upstream of its calcium binding C2 domain, p47^phox^ is cleaved in the middle of its PX (phox) domain (74,75). The PX domain of the p47^phox^ subunit of NADPH oxidase, once phosphorylated, mediates the recruitment of this subunit from the cytosol to the phagosome membrane via binding to acidic phospholipids (75). Simultaneously, V8 substrate Rac2 will become activated via GDP to GTP conversion, translocate to the membrane and interact with the NADPH subunits, forming a fully active phagocyte oxidase complex (6). To further emphasize the importance of Rac2, studies have shown that activation of the NADPH complex after FcyR mediated phagocytosis is completely dependent on this protein in murine neutrophils (76). As mentioned, Rac2 is a multifunctional protein also displaying a role in leukocyte migration, however in this regard, targeting of Rac2 by V8 could promote *S. aureus* pathogenesis as a consequence of abrogated ROS production. This is supported by evidence showing patients with decreased levels of this protein experience an autosomal dominant immunodeficiency syndrome characterized by defects in overall neutrophil functionality (72,77). This syndrome is seen mainly in infants where their neutrophils have decreased ROS levels, dampened chemotaxis, polarization and azurophilic granule secretion (77). Moreover, patients with Rac2 deficiency present with severe and recurring bacterial infections, accompanied with poor wound healing similar not only to LAD-1, but chronic granulomatous disease (CGD) (70,77–78).

Further evidence suggesting V8 is involved in abrogation of ROS production comes from our observed cleavage of the calcium binding chaperone calreticulin. This versatile protein, found to reside not only in lumen of the endoplasmic reticulum, but also on the neutrophil cell surface, has emerged as an important immune response regulator involved in ROS production and pathogen clearance (79–81). In support of this, previous work investigating the activity of a synthetic antimicrobial peptide demonstrated its ability to bind calreticulin on the surface of neutrophils and induce neutrophil activation and enhance superoxide production via GPCR signaling (80). Consequently, this peptide exhibited significant anti-staphylococcal action in MRSA infected mice (82). While this AMP may not be in circulation in the normal host environment, this previous study presents evidence for calreticulin as a binding target to induce neutrophil activation, a phenomenon that can be potentially dampened due to V8 cleavage. Calreticulin possess three important domains including the N, P and C-domain, with the former two involved in chaperone function and the C-domain involved in calcium binding (79–81). We show V8 cleavage of this protein after glutamic acid residues at positions 128 and 342 (**Table S2**), corresponding to both the N- and C-domains, likely abrogating their function.

Intriguingly, additional studies on this molecule have highlighted its pivotal role in protein synthesis and maturation of the ROS mediator, myeloperoxidase, emphasizing the paramountcy of calreticulin in the innate immune response (79). Proposed to stabilize the immature form of MPO to facilitate heme binding (79), cleavage of calreticulin by V8 points would engender significant dysregulation of ROS production. As such, our data establishes multiple routes by which V8 can extinguish intracellular neutrophil defense mechanisms, leading to prolonged survival and persistence within the host.

In support of this, we validated V8 cleavage of ROS mediator, pyruvate kinase PKM2. This protein was found to be cleaved three times by V8, in the barrel domain at positions 355 and 283, with the latter being just upstream of the active site or bait region, and the C-terminus at position 481 (**Figure 4D**). This observed cleavage pattern could give rise to four different degradation products as outlined in the protein schematic (**Figure 4D**). While we observed a decrease in intensity of the full length PKM2 protein and modified form in the V8 treated lanes, we also see an additional band between ∼25-35kDa, that corresponds to the molecular weight of cleavage fragments predicted by our N-terminomic data (**Figure 4D**).

### N-terminomics predicts V8 modulation of phagocytosis

Our study also recorded V8 cleavage of many important neutrophil structural proteins, including myosin-9, tubulin chains, filamin A, and actin subunits. These cytoskeletal proteins are responsible for maintaining important neutrophil functions and characteristics, such as motility, cell shape, membrane organization, and phagocytosis (83,84). Previous work has shown that *S. aureus* can prevent phagocytosis via anti-opsonic strategies, such as cleavage of complement C3 by aureolysin, or via the binding of immunoglobulins by either staphylococcal protein A (Spa) or staphylococcal binding of immunoglobulins (Sbi) protein (3). Herein, our data provides an additional route for how *S. aureus* can modulate this host defense tactic facilitating immune evasion. Defects in phagosomal cup formation and thus phagocytosis could arise from V8 cleavage of myosin-9, or non-muscle myosin heavy chain IIA, an essential motor protein driving actin polymerization and subsequent extension of neutrophil pseudopod protrusions (84). This notion is supported by a study showing that in the presence of blebbistatin, a specific inhibitor of myosin II, neutrophil phagocytosis is abrogated (85). Given that V8 also cleaves various actin proteins, this suggests dysregulation of the entire process of phagocytosis. Furthermore, as pseudopod extension and actin assembly also propel neutrophils during migration, V8 proteolysis of these proteins provide an additional layer of evidence suggesting that this protease can impede neutrophil migration. Once again, this is confirmed by previous work that highlights how application of a myosin IIA inhibitor limits not only phagocytosis but neutrophil migration (85).

We took a different approach to validation, whereby we subjected purified human beta actin protein to digestion by V8, with the resulting banding patterns visualized by Coomassie blue staining (**Figure 5A**). Our N-terminomic data captured cleavage of this molecule at positions 5, 148, 227 and 317. The purified protein used was a His-tagged truncated variant with a molecular weight of ∼35kDa; the predicted banding patterns and their molecular weights are outlined in the protein schematic. Following V8 proteolysis, we observed two fragments at ∼12kDa and ∼15kDa, directly corresponding to our N-terminomic results.

**Figure 5.**
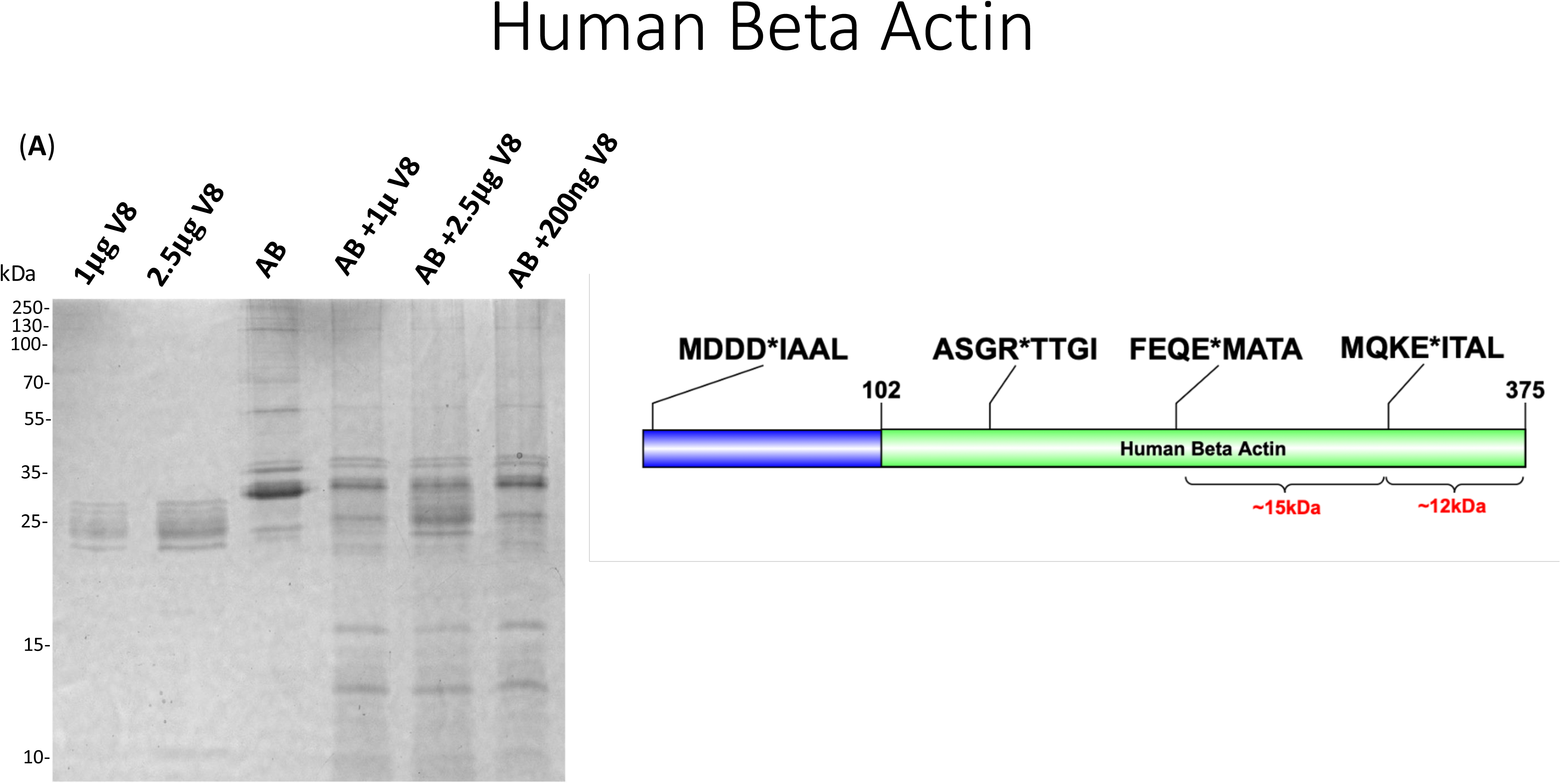
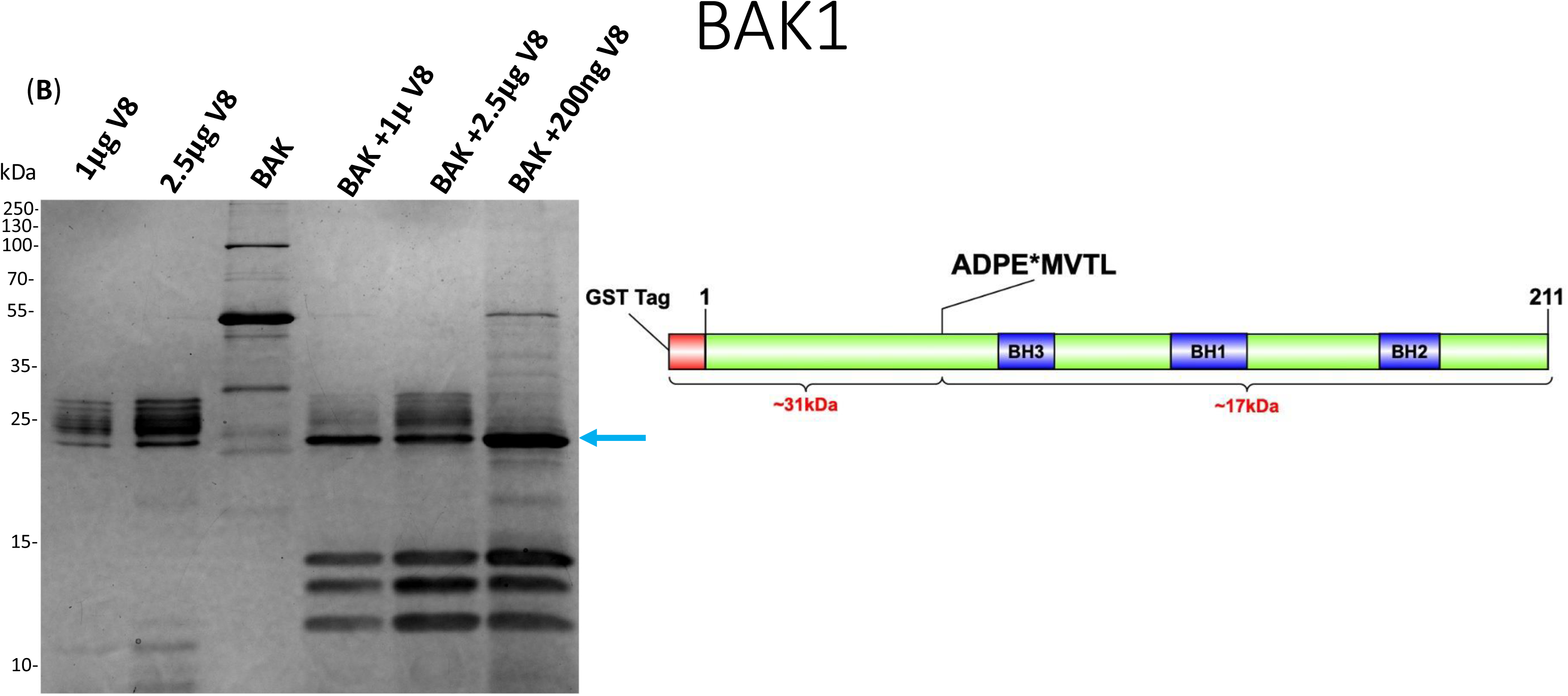
SDS-Page validation of neutrophil V8 targets using purified recombinant proteins. Recombinant proteins of (**A**) Human beta actin and (**B**) BAK1 were treated with varying concentrations of the V8 protease and compared to untreated controls. Purified V8 was also included as a control. A schematic of the corresponding protein is presented adjacent to the Coomassie Blue stained gel. Green coloring indicates the corresponding portion of the protein that was purchased. Cleavage sites detected by TAGS-CR are highlighted with the asterisk and the surrounding residues, as well as the resulting degradation products and their corresponding molecular weights. The blue arrow indicates bands that were excised for in-gel digest for (**B**) BAK1.

### V8 degrades apoptotic signaling proteins

TAGS-CR captured V8 cleavage of pro-apoptotic proteins, including the BH3-interacting domain death agonist (BID) and apoptosis regulator BAK1, and its homolog BAX. These V8 targets are all pro-apoptotic members of the Bcl-2 family that together are essential gateway molecules for inducing apoptosis (86). BH-3 only proteins, such as BID, BIM, BAD and PUMA, drive the apoptotic cascade as they induce oligomerization of BAK1/BAX, resulting in mitochondrial outer membrane permeabilization (MOMP), followed by the release of cytochrome c and subsequent activation of downstream caspases, which function as major mediators of programmed cell death (86). In healthy cells, the response to apoptotic stress begins with cleavage of BID by caspase-8 at position 59 to form a truncated version of the molecule, tBID, known to induce BAK1/BAX association and subsequent MOMP formation (87). Given this, our data points to V8 behaving as a potential activator of the apoptotic process, due to V8 cleavage of BID at position 53, potentially mimicking the caspase-8 effect.

Beyond BID, the apoptotic cascade remains complex as the mechanisms of BAK1/BAX activation are not well understood (88). However, these molecules have been found to necessitate structural reconfiguration prior to pore formation, including N-terminal conformation changes (89–91). As such, previous work has shown that dissociation of the BAK1 α1 helix is necessary for unfolding and exposure of its hydrophobic binding pocket between α2-α5, which mediates interaction with BH-3 only activators such as tBID (92). From our data V8 cleaves BAK at position 60, found in the α1-α2 loop, which would result in release of the N-terminal segment and α1 helix (92). Additional studies have explored influence of the α1-α2 loop on BAK1 activity and determined that it elicits a suppressive effect on BAK1 activation via stabilization of the molecule (93). Interestingly, it was found that amino acid M60 in the α1-α2 loop was responsible for stabilization effect of this structure, ultimately preventing downstream activation and subsequent apoptotic pore formation (93). Taken together, V8 cleavage within the α1-α2 loop at position 60 indicates that this protease may trigger activation of the apoptotic cascade.

While structural rearrangements in BAK1 are akin to those in BAX, it has been hypothesized that α1 dissociation is also necessary for BAX activation (92). Herein, we see that V8 may also elicit a similar effect on BAX, as observed for BAK1, cleaving at position 45 in the same structural domain, the α1-α2 loop (95). It is well documented that *S. aureus* can strategically manipulate host apoptotic processes to permit survival and persistence within the human host. Promotion of apoptosis by virulence factors, such as α-toxin or the pore-forming leukocidins, has been shown to not only induce tissue injury and exacerbate inflammation, but allow for escape and spread to new host niches (96). Given that the secreted proteases have been recognized for their role in promoting dissemination during infection, our study highlights the possible contributory effect of V8 to this phenomenon via cleavage of BID, BAK1 and BAX.

As evidence for this claim, we validated cleavage of BAK1. Recombinant GST tagged BAK1 was digested with purified V8, however there was a discrepancy between predicted and corresponding molecular weights of the resulting degradation products. Therefore, we excised the three bands in line with the blue arrow across the different V8 treatment conditions (**Figure 5B**) and subjected them to in-gel tryptic digest and subsequent MS/MS analysis. In addition to possible self-degradation of the V8 enzyme, the 26kDa GST tag may also have undergone digestion by this protease, therefore we included both protein sequences for spectral assignment when searching the raw data. We considered peptides identified in at least two out of the three treatment conditions as well as having the presence of Glu or Asp residues at either the N- or C-terminus, as constituting V8 semi-tryptic peptides (**Table S4**). From this we captured 5 reliably detected peptides, 4 identified as the GST tag and 1 corresponding to the BAK1 protein with a start position of 60 as captured by TAGS-CR.

### The V8 protease facilitates survival of *S. aureus* within human leukocytes

Given that our N-terminomic data suggests clear disruption of proper neutrophil functioning by the V8 protease, we sought to assess whether this enzyme contributes to intracellular survival of *S. aureus* in human neutrophils. As such, a *S. aureus* Δ*sspA* mutant was compared to the USA300 LAC wild type for viability in a model of neutrophil survival. At 24h post-infection we determined a clear decrease in the survival of the mutant strain, with bacterial burden decreased around two-fold in comparison to WT (**Figure 6**). As such, loss of the V8 protease negatively impacts survival of *S. aureus* inside the phagolysosome of human leukocytes as would be expected from our N-terminomic data. Of note, a complementing strain was also included in this experimental model (data not shown), however it failed to complement due to instability of the plasmid; something previously noted in other studies (97–99).

**Figure 6.**
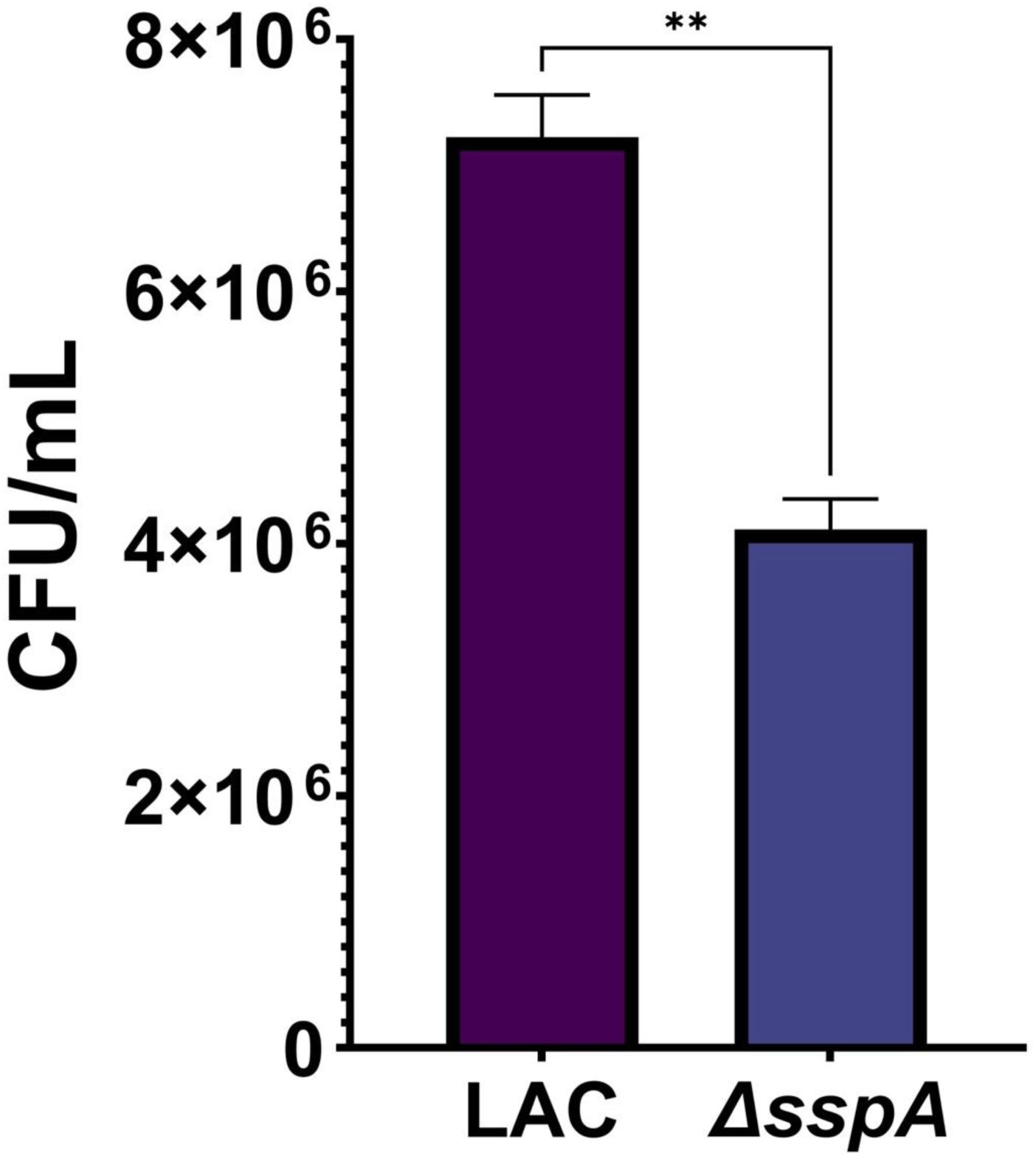
The V8 protease promotes intracellular survival of *S. aureus* within human leukocytes. HL-60 derived human neutrophils were infected with either the *S. aureus* LAC wild type or the *sspA* (V8 encoding gene) mutant strain using an MOI of 13.5. CFU/mL was then determined after a 24-hour period and data displayed represents the average of 3 biological replicates for each strain. The significance of relative bacterial burden between strains was determined using an unpaired t-test with unequal variance, **p=<0.01. Error bars are ± SEM.

## SUMMARY AND CONCLUSION

The function of proteases as enzymes is defined by their action on other proteins – thus, to understand the role of a protease, one must identify its substrates (25–33, 101). A common misconception is that proteolysis leads to complete degradation. Instead, proteases can cleave with exquisite specificity (particularly V8), rather than randomly, recognizing substrates via key motifs (37, 102–107). Our ability to capture such events has been slow as conventional strategies to assess proteolysis are low throughput, and involve one at a time approaches. However, recent advances in mass spectrometry have led to the field of degradomics, allowing us to capture proteolysis on a global scale (33–36). Herein, we present a novel N-terminomic approach – TAGS- CR - that facilitates the rapid, efficient and streamlined resolution of proteolytic events in complex samples.

As neutrophils remain steadfast as the front-line defense against invading *S. aureus*, it is pivotal to understand the molecular mechanisms employed by this pathogen to thwart immune responses and foster infection (52). Using TAGS-CR we determined that V8 can potentially dysregulate a wide breath of neutrophil functions via targeted cleavage of ∼350 host proteins. Specifically, we captured proteolysis of critical factors involved in neutrophil adhesion and migration, a fundamental process necessary for pathogen clearance and infection resolution (22). We also showed that V8 can disrupt phagolysosomal defense and apoptosis serving to facilitate *S. aureus* survival and persistence within the host (12,22,96). Indeed, in the absence of *sspA* (the V8 encoding gene)*, S. aureus* has an impaired ability to survive during engagement with human neutrophils.

We acknowledge that our experimental design may have perceived limitations in the form of access of protease to substrate, facilitated by our use of whole, lysed neutrophil proteomes. We counter this contention with the notion that *S. aureus* is well adapted to intracellular survival within neutrophils, not only escaping the lethal effects of the phagolysosome, but also thriving within the cytoplasm of these immune cells (108). As such, *S. aureus* and its proteases would readily have access to most if not all intracellular machinery within neutrophils. In addition, the secreted proteases function in concert with other secreted virulence factors of *S. aureus* (26). Consequently, lytic action on neutrophils by the leukocidins would again readily provide access for the proteases to intracellular components of these immune cells.

As such, we suggest that our work illustrates application of a unique N-terminomic strategy and demonstrates its potential in providing unparalleled insight into *S. aureus* host-pathogen interaction. We present the detailed mapping of the human neutrophil pathodegradome, which not only propels bacterial protease substrate identification but shines new light on the immunomodulatory mechanisms implemented by *S. aureus.* Furthermore, as neutrophil dysfunction often results in recurrent *S. aureus* infections, this further emphasizes the significance of our results and offers insight that could facilitate the development of effective treatments and preventative strategies.

## Acknowledgements

This study was supported by grants AI124458 and AI157506 (L.N.S.) from the National Institute of Allergy and Infectious Diseases.

## METHODS

### Neutrophil Proteome V8 treatment

Six flasks of human HL-60 leukemia cells were grown at 37°C with 5% CO_2_ in RPMI supplemented with 10% fetal bovine serum (FBS) and penicillin and streptomycin. HL-60 cells were differentiated into neutrophil-like cells via stimulation with 1.25% DMSO. Differentiation was allowed to proceed for 5 days and then cells were centrifuged at 1,000g for 5 minutes. Cell pellets were washed twice with PBS prior to protein extraction. A 50µl portion of each wet cell pellet was resuspended in 500µl of the Mammalian Protein Extraction Reagent M-Per (Thermofisher Scientific). Proteins were extracted by vigorous shaking for 20 minutes, pooled and then quantified using a Pierce 660nm protein assay (ThermoFisher Scientific). Neutrophil proteomes were standardized to 1mg/ml and incubated ±200ng of V8 protease (MilliporeSigma), at 37°C for 16h. Following incubation, protease activity was quenched with dry guanidium hydrochloride (6M final concentration) prior to N-terminomic experimentation.

### N-terminomics: Terminal Amine Guanidination of Substrates (TAGS)

Following protease treatment, samples were incubated with 20mM DTT at 95°C for 10 minutes and centrifuged at 17,0000g to remove remaining non-solubilized proteins. Iodoacetamide was added to a final concentration of 40mM and incubated in the dark at RT for 30 minutes. Alkylation was quenched with 40mM DTT. Guanidination of protein N-termini was performed by incubating samples with 500mM of triethyl ammonium bicarbonate (TEAB) and 500mM of 1H-pyrazole-1-carboxamidine hydrochloride (HPCA) at 95°C for 10 minutes. Samples were then quickly chilled on ice for 5 minutes. In accordance with the S-trap protocol (PROTIFI), phosphoric acid was added to a final concentration of 1.2 %. In a 1:7 volumetric ratio, S-trap buffer (10% v/v TEAB pH 7.5, 90% v/v MeOH) was mixed with each sample. Proteins were then bound to the S-trap midi column matrix by repeated flow through at 4,000g for 1 minute. Samples were washed a total of three times, each by adding 3ml S-trap buffer and centrifuging as above. For protein digestion, Trypsin/P was added at 1:100 enzyme:protein (10µg of trypsin in 50mM TEAB pH 8.1) and samples were incubated at 37°C for 18h. Elution of peptides was performed according to the S-trap protocol, with a series of centrifugations starting with 50mM TEAB pH 8.1, then 0.2% (v/v) formic acid and finally with 50% acetonitrile. In a vacuum centrifuge, samples were evaporated to ∼1/3 of their final volume. A three-step incubation process was then performed to sulfonate peptides to facilitate downstream enrichment (Lai et al., 2015). Samples were first incubated with 100mM HEPES pH 7.5 and 20mM FBDA and 20mM SCBH for 1h at RT. This was followed by 2 additional incubations with 20mM FBDA and 20mM SCBH at RT, the first for 1h and then for 16h. FBDA and SCBH were made fresh prior to each incubation, with the final concentration in the reaction being 60mM FBDA and 60mM SCBH. Following final incubation, excess FBDA and SCBH was quenched by the addition of 2M Tris (pH 8) to a final concentration of 100mM. Using Sep Pak C18 columns (Waters), samples were desalted according to the manufacturer’s protocol. Using a vacuum centrifuge, samples were dried to completion. Strong cation exchange was performed using mini pierce strong cation exchange spin columns (ThermoFisher Scientific). Briefly, peptides were resuspended in 300µl of SCX buffer A (5mM Ammonium Formate, pH 3.0, 25% (v/v) CAN) and then vortexed and sonicated for 5 minutes each. SCX columns were equilibrated by centrifuging 400µl of SCX buffer A for 5 minutes at 2,000g (all SCX centrifugations took place using these parameters unless stated otherwise). Samples were then added to the SCX column and centrifuged, followed by two washes with the addition of 400µl of SCX buffer A and centrifugation. Peptides were eluted from the SCX column with the addition of 200µl of SCX buffer B (500mM Ammonium Formate, pH 6.0, 25% (v/v) ACN) followed by a 1-minute incubation and then centrifugation. This was repeated once to bring the final volume to 400ul. Accordingly, to reduce the ACN to 7.25%(v/v), 1.2mL HPLC water was added to each tube. Samples were acidified with the addition of 80µl of trifluoroacetic acid and an additional desalt was performed using Sep Pak C18 columns (Waters) according to manufacturer’s instructions. Finally, peptides were dried to completion using a vacuum centrifuge and stored at 4°C prior to mass spectrometry.

### Mass Spectrometry

The N-terminally enriched peptides obtained following TAGS-CR were resuspended in 0.1% formic acid. Aliquots of 5µl were separated on a 50cm Acclaim PepMap 100 C18 reversed-phase high-pressure liquid chromatography (HPLC) column (Thermo Fisher Scientific) using an Ultimate3000 UHPLC (Thermo Fisher Scientific) with a 120 min gradient (2%-32% aceto-nitrile with 0.1% formic acid). A hybrid Quadrupole-Orbitrap instrument (Q Exactive Plus; Thermo Fisher Scientific) analyzed peptides using data dependent acquisition, with the top 10 most abundant ions being selected for MS2 analysis.

### Data Analysis

Raw files were processed using MaxQuant (100). The andromeda search engine included in this program was used to assign MS2 spectra to the reference human proteome (UniProt ID UP000005640). For this N-terminomics experiment, a semi specific digestion setting with trypsin/P was used. Fixed modifications included Carbamidomethyl (C), Guanidination (K), while variable modifications included Oxidation (M), Acetyl (Protein N-term) and Guanidination (N-term). Each experiment was injected into the mass spectrometer twice to give n=6 for both control and V8 treated conditions and labelled appropriately in the MaxQuant software. The match between runs feature was also selected and peptide and protein FDR were both set to 1%. The resulting modified peptide output file was analyzed using R programming language whereby contaminants and unmodified peptides were removed and peptides considered for analysis were identified in at least two of the three separately conducted experiments. Subset lists of peptides, proteins, N-termini, and cleavage sites were generated to allow for a binary comparison of unique cleavage events between ±V8 conditions.

### SDS-PAGE and Immunoblotting

SDS-PAGE was performed using either 12% or 15% self-cast gels, or 4-20% gradient gels (BioRad). Coomassie blue was used to stain gels with recombinant proteins, while gels for digested neutrophil proteomes were immediately transferred to a PVDF membrane at 20 Volts for 45 minutes. Following transfer, membranes were blocked (5% milk in Tris buffered saline–Tween 20) for 1h at RT with rocking. Membranes were then exposed to primary antibody (1:1300 – 1:1000 dilution of antibody to blocking buffer) for 16 hours at 4°C and washed 3 times for 10 minutes with blocking buffer. Blots were exposed to the secondary antibody anti-rabbit IgG-HRP conjugate (Cell Signaling Technologies) at a 1:5000 dilution of antibody to blocking buffer for 1h at RT with rocking, and then washed 3 times for 10 minutes with blocking buffer. The Supersignal West Pico chemiluminescent substrate (Thermo Fisher) was added to the blot prior to imaging. Blots were captured using developing film having been exposed from 15 minutes to 18h. Primary antibodies used included: Integrin alpha-L (abcam, ab52895), Kindlin-3 (Thermo Fisher Scientific, #PA5-30847), Myeloperoxidase (abcam, ab134132), Pyruvate kinase PKM2 (Thermo Fisher Scientific, #PA5-28700). Purified recombinant proteins used included: Human beta actin (abcam, ab240844) and Human BAK1 (Abnova, 89-936-369).

### In-Gel Digestion

Designated V8 treated BAK1 bands were excised and cut into 1mmx1mm cubes. Samples were washed twice with 200µl of 50/50 ACN/TEAB by vortexing for 15 minutes, discarding the wash each time. Gel pieces were then dehydrated by covering with ACN for 10 minutes. Once ACN was removed, samples were rehydrated with 50µl of 100mM TEAB and left to incubate for 5 minutes. To each sample, an equal volume of ACN was then added to achieve a 1:1 ratio of ACN:100mM TEAB and a final wash was performed. Following removal of the wash, samples were dried in a vacuum centrifuge for 5 minutes and then incubated at 55°C for 30 minutes in the presence of 100µl of 50mM DTT. Once samples were allowed to return to RT, they were then incubated in 100µl of 100mM IAA in the dark at RT. IAA was removed and gel pieces were washed three times as described above and then dried for 5 minutes in a vacuum centrifuge. To each sample, 100ng of trypsin was added, followed by overnight incubation at 37°C. Peptides were collected by transferring the supernatant to a fresh microcentrifuge tube and by performing two subsequent washes, adding each to the new designated sample tube. Once peptides were dried to completion in a vacuum centrifuge, mass spectrometry was performed as described above and raw files were processed using MaxQuant (100). The resulting peptide file was analyzed and contaminants, peptides not found in at least 2/3 BAK1 V8 treated lanes, as well as non-V8 semi-tryptic peptides, were removed.

### Data Deposition

Raw mass spectrometry data for this study was deposited as two submissions to the ProteomeXchange Consortium using the PRIDE repository with the dataset IDs PXD057551 and PXD057579 for the neutrophil N-terminomics and the BAK1 in-gel digestion, respectively.

